# Gut microbiota-mediated conversion of mangiferin to norathyriol alters short chain fatty acid and urate metabolism

**DOI:** 10.1101/2024.12.19.629397

**Authors:** Daan Bunt, Markus Schwalbe, Fittree Hayeeawaema, Sahar El Aidy

**Affiliations:** Host-Microbe Interaction, Groningen Biomolecular Sciences and Biotechnology Institute (GBB), University of Groningen, 9747 AG Groningen, The Netherlands; Microbiome Engineering, Microbiology Department, Swammerdam Institute for Life Sciences (SILS), University of Amsterdam; Stratingh Institute for Chemistry, University of Groningen, Nijenborgh 7, 9747 AG Groningen, The Netherlands

**Keywords:** mangiferin, norathyriol, C-glycosides, metabolites, gut microbiota, urate, SCFAs, sequencing, fermentation

## Abstract

Mangiferin (MAN), a natural C-glycosylxanthone, is recognized for its health-promoting effects in traditional medicinal preparations. However, its poor bioavailability and limited intestinal permeability restrict its direct biological activity *in vivo*. Previous studies have suggested a potential bacterial breakdown of MAN into norathyriol (NOR), an aglycone with significantly higher bioavailability and absorption. Yet, the prevalence of MAN-metabolizing microbes, the presence of MAN or NOR within the gut microbial community, and their effects on the composition and metabolic activity of the gut microbiome remain unclear. In this study, fecal samples from healthy adult volunteers treated with MAN revealed its conversion to NOR, with interindividual variation attributed to the uncultured bacterial strain CAKRHR01 *sp934339005*. While MAN had minimal impact on microbial composition and metabolic activity, NOR treatment significantly increased pH, reduced overall bacterial cell counts, and selectively suppressed short-chain fatty acid-producing bacteria, including *Faecalibacterium prausnitzii* as well as urate consumers, such as *Enterocloster bolteae*. These findings underscore the potential of NOR to modulate gut microbial activity and highlight the importance of understanding microbiome-mediated metabolism when assessing the health implications of phytochemicals.

## Introduction

Mangiferin (MAN) is a natural C-glycosylxanthone predominantly found in various plant species, most notably, in the leaves, stems, bark, and fruits of *Mangifera indica* (mango tree). These plant parts are widely used in traditional medicine, particularly across the Indian subcontinent, to treat various ailments, including gastrointestinal disorders, bronchitis, and diabetes.^1^ Studies on isolated MAN have demonstrated a broad spectrum of biological activities, including anti-diabetic, gastroprotective, neuroprotective, and immunomodulatory effects.^2^ Consequently, MAN is considered a critical bioactive component in traditional medicinal preparations derived from *Mangifera indica*.

Despite its promising bioactivity, studies have demonstrated that MAN exhibits negligible oral bioavailability in rats, estimated at approximately 1-2%.^3–5^ This poor bioavailability is primarily attributed to its poor solubility and permeability within the intestinal tract.^6^ Notably, even at relatively high oral doses ranging from 50-500 mg/kg, MAN remains undetectable in rat plasma.^7,8^ Such pharmacokinetic limitations are characteristic of compounds with C-glycosidic bonds, which typically exhibit low absorption and bioavailability.^9^ Additionally, the metabolic resistance of the C-glycosidic bond due to their stability under acidic conditions and resistance to cleavage by glycosidases, distinguishes them from O-glycosidic bonds, which are readily hydrolyzed by acidic environments or glycosidases of human and bacterial origin.

Although C-glycosides are generally more metabolically stable, evidence suggests that bacterial enzymes, can deglycosylate them.^10–12^ This deglycosylation process is distinct from typical glycosidase activity and involves a two-step mechanism: oxidation of the 3-hydroxyl group on the glycosyl moiety, followed by elimination of the aglycone from the resulting 3-oxo-glycosyl moiety **(Figure 1A)**.^12,13^

**Figure 1:**
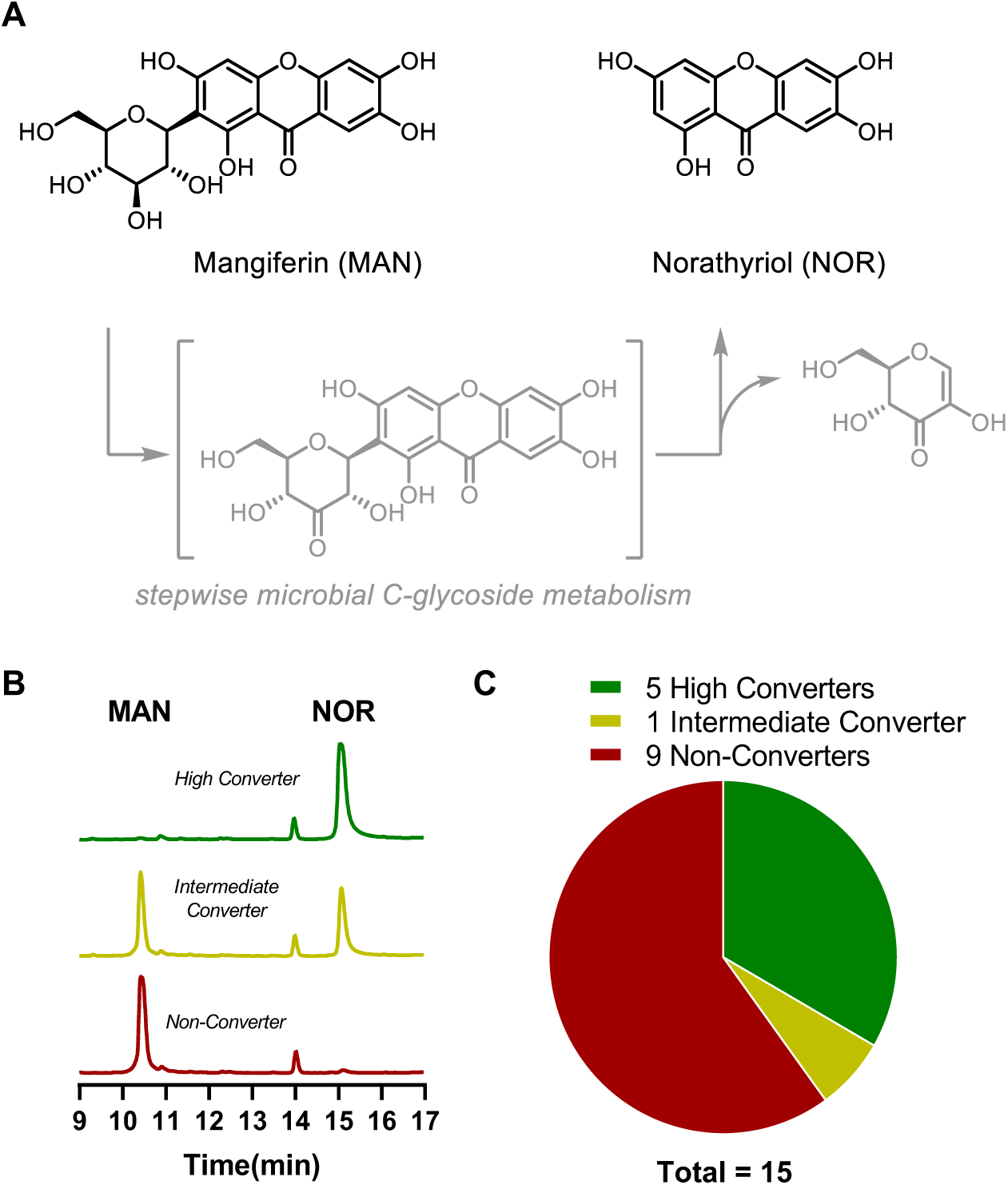
Inter-individual variability in the microbial conversion of mangiferin (MAN) to norathyriol (NOR) by the gut microbiota. (A) Chemical structures of MAN and NOR, highlighting the stepwise microbial deglycosylation pathway. (B) Representative HPLC-UV chromatograms illustrating a high converter, partial converter, and non-converter. (C) Pie chart depicting the distribution of conversion types among tested fecal samples (n = 15).

In contrast to MAN, its deglycosylation product, norathyriol (NOR), exhibits significantly higher oral bioavailability in rats, estimated at 30.4%, despite undergoing hepatic metabolism.^14^ A distinctive double-peak phenomenon in the concentration of NOR has been observed in the portal vein, indicative of its enterohepatic circulation.^15^ This prolonged intestinal exposure to NOR, coupled with its enhanced bioavailability, underscores its greater potential for exerting biological effects in vivo compared to MAN.

Given the distinct pharmacokinetic and biological activities of MAN and NOR, this study employs an integrative approach that combines *in vitro* fermentation, targeted and untargeted metabolomic analysis, and deep metagenomic sequencing. Through these complex interactions, this study unravels the mechanisms underpinning the metabolic fate of MAN and NOR in the gut. The findings highlight their potential role in shaping gut microbial ecology and its interactions with intestinal tissue, offering a deeper understanding of how gut microbiota modulate the bioactivity of these dietary compounds.

## Material and Methods

### Chemicals

Mangiferin (M2886) was acquired from TCI Europe. Norathyriol was prepared according to a literature procedure.^16^

### Anaerobic mangiferin fermentation experiments

Fecal samples were collected from healthy volunteers, who provided written consent for participation, and stored immediately in liquid Amies at -80°C.^17^ Precultures were prepared by inoculation of 50 mg homogenized fecal sample into 3 mL Gifu Anaerobic Broth (GAM, HiMedia Laboratories) medium, prepared according to the manufacturer’s instructions, and incubating under anaerobic conditions (5% H_2_, 5% CO2, balanced with N_2_) in a Coy Laboratory Anaerobic Chamber (neo-Lab Migge GmbH, Heidelberg, Germany), at 37 °C for 24 h. A 10% inoculum from the preculture was then re-inoculated into 3 mL fresh GAM at 37 °C for an additional 24 h. This process was repeated using 1% of the second culture. Subsequently, 60 µL samples from the cultures were centrifuged, and the cell pellets were resuspended in 3 mL carbohydrate-free Enriched Beef Broth (EBB).^18^ To this, 150 µL of a 500 µM MAN solution in DMSO was added. The mixture was incubated at 37 °C for 48 h. After the incubation, a 250 µL aliquot was collected, immediately mixed with cold methanol, and stored at -20 °C until analysis by HPLC.

### Aerobic mangiferin fermentation experiments

Precultures were prepared from ileostomy samples, which were previously collected from otherwise healthy patients and published in our previous study^19^, were prepared by inoculating glycerol stocks in 3 mL enriched beef broth (EBB) growth medium^18^ at 37 °C for 24 h. A 10% of the inoculum from the pre-culture was re-inoculated into 3 mL fresh EBB and incubated at 37 °C for another 24 h. The process was repeated using 1% of the second culture. From the cultures, 60 µL samples were centrifuged, the cell pellets were resuspended in 3 mL of either EBB or carbohydrate-free EBB medium. To each sample, 150 µL of a 500 µM MAN solution in DMSO was added. The mixture was incubated at 37 °C for 48 h. After incubation, a 250 µL aliquot was collected, mixed with cold methanol, and stored at -20 °C until analysis by HPLC.

### Fermentation experiment using SIFR® technology

Fecal samples were collected from healthy adult human volunteers (*n* = 6), in accordance with the ethical guidelines approved by Ethics Committee of the University Hospital Ghent (reference number BC-09977). The selection criteria for the donors included: age 18-65, no antibiotic use in the past 3 months, no gastro-intestinal disorders (cancer, ulcers, IBD), no use of probiotic, non-smoking, alcohol consumption < 3 units/d and BMI < 30.

The study consisted of 3 treatment arms: 500 µM MAN, 500 µM NOR, and no-substrate control (NSC), with each condition run in technical triplicate per donor. Individual bioreactors were processed in parallel in a bioreactor management device (Cryptobiotix, Ghent, Belgium). Each bioreactor contained 5 mL of nutritional medium (M0017, Cryptobiotix, Ghent, Belgium) containing 500 µM MAN, 500 µM NOR, or no substrate, and fecal inoculum of the healthy donors. The mixtures were incubated at 37 °C for 24 h, simulating colonic fermentation. Analysis was conducted at two timepoints: 0 hours (for the NSC condition) and 24 hours (for all conditions). At both timepoints, pH and gas production were measured, and cell counts were determined using flow cytometry. The concentrations of acetate, propionate, butyrate, valerate and branched-chain fatty acids (sum of isobutyrate, isovalerate, and isocaproate) were quantified using GC-FID.^20^ Cell pellets were obtained by centrifuging 1 mL of culture and stored at -80 °C until further processing. Filter-sterilized supernatant (1 mL) and raw sample (0.5 mL) were collected and kept at -80 °C until further analysis.

### DNA isolation and Library preparation

The DNA was extracted using Tiangen Magnetic Soil And Stool DNA Kit. Library preparation was carried out using the Novogene NGS DNA Library Prep Set (Cat No.PT004) with index codes assigned to each sample. Briefly, genomic DNA was randomly fragmented to size of 350bp. The DNA fragments were end-polished, A-tailed, ligated with adapters. Size selection was performed, followed by Rolling Circle Amplification. The resulting PCR products were purified using the AMPure XP system and analysed for size distribution using an Agilent 2100 Bioanalyzer (Agilent Technologies, CA, USA). Quantification was performed using real-time PCR. The library was then sequenced using the Novaseq X plus flow cell with PE150 strategy. All sequencing data is available under the following project accession PRJNA1199756.

### Bioinformatic analysis

Briefly, raw reads were trimmed with trimGalore, and host DNA was removed with Hocort^21^ in biobloom mode, yielding “clean reads”. The clean reads were subsequently assembled using both SPAdes^22^ in meta mode or MEGAHIT ^23^ in parallel. Assemblies were subjected to binning using BASALT.^24^ The resulting bins were dereplicated using dRep^25^, yielding a non-redundant bin set. Taxonomic annotation of bins was performed using GTDB-Tk.^26^ In parallel, gene prediction was carried out on all (redundant) bins using Prokka^27^ and the predicted proteins were aggregated into a non-redundant protein catalogue using MMSeqs2.^28^ Bin abundances were determined by mapping clean reads against the non-redundant bin set using Bowtie 2^29^ and mapped reads per contig were obtained by SAMtools idxstat.^30^ These steps were implemented in a Nextflow pipeline^31^, with environment files showing respective tool versions available online (https://github.com/m4rku5-5/metagenomicsPipeline.git). Read counts were converted to pseudo TPM values using the formula: pTPM = mapped reads / (total bin length / 1000) / (Total reads in sample / 1000000).

To generate absolute abundances for each bin, pTPM values were multiplied by cell counts (determined by flow cytometry, see above) and these values were used for downstream analysis. Compositional analysis was performed using the statistical programming language R (v4.3.3), with the following packages: phyloseq (v1.46.0), microViz (v0.12.1), vegan (v2.6), ggpubr (v0.6.0) and tidyverse (v2.0.0). Differential abundance of features was assessed using Maaslin2^32^ (v1.16.0), including the Donor individual as random effect.

### Analysis of DfgABCDE and uric acid gene cluster

Sequences were obtained from NCBI and searched against the metagenomic non-redundant protein catalogue using diamond^33^ in ultra-sensitive mode. Only sequences with more than 30% sequence identity and 80% query coverage were retained. Proteins were made non redundant using their cluster membership and then compared to their respective taxonomy assignment. Genomic arrangements were visualized from their corresponding GFF files using the gggenes package (v0.5.1).

### HPLC-UV analysis and sample preparation

Protein precipitates were removed by centrifugation at 20,000 × g for 10 min at 4 °C. The supernatant was transferred to a new tube, and the methanol fraction was evaporated in a Savant speed-vacuum dryer (SPD131, Fisher Scientific) at 45°C for 2 h. The aqueous fraction was reconstituted to 1 mL with 0.7% HClO_4_. For anaerobic incubation samples, these were filtered and injected into the HPLC-UV system (Waters 2695 Alliance Separations module; TSP UV6000 detector measuring at 256 nm). The analysis was performed using a C18 column (Phenomenex Kinetex 5 μm, C18 100 Å, 250 × 4.6 mm) with a gradient of water/methanol with 0.1% formic acid (0-13 min, 90% to 30% H_2_O; 13-15 min, 30% to 0% H_2_O; 15-16 min, 0% to 50% H_2_O; 16-18 min, 50% to 90% H_2_O) and a flowrate of 1 mL/min. The column temperature was maintained at 35 °C. Data recording and analysis were performed using Chromeleon software (version 6.8 SR13) and GraphPad Prism 8. For aerobic incubation and SIFR**®** experiments, were filtered and injected into a UHPLC system (Dionex UltiMate 3000 autosampler; Dionex AltiMate 3000 LPG-3400SD pump, Thermo Fisher Scientific, Waltham, Massachusetts, USA). Analysis was performed on a C18 column (Phenomenex

Kinetex 5 μm, C18 100 Å, 250 × 4.6 mm) with a gradient of water/methanol with 0.1% formic acid (0-7 min, 50% to 5% H_2_O; 7-8 min, 5% to 50% H_2_O; 8-9 min, 50% H_2_O) and a flow rate of 1 mL/min. The column temperature was set to 40 °C. Data recording and analysis were performed using Chromeleon software (version 6.8 SR13) and GraphPad Prism 8.

### Untargeted metabolomics

To 50 ul of sample 250 ul of ice-cold MeOH were added, followed by thorough mixing and sonication in a chilled sonication bath. Subsequently 500 ul of ice-cold CHCl3 were added, mixed for 2 minutes and incubated on ice for 10 minutes. To this 200 ul of cold water was added, followed by mixing for 2 minutes, incubation on ice for 10 minutes and centrifugation at 1000 rpm for 5 min @ 4 °C. Subsequently 400 ul of the upper (polar) and lower (apolar) phases were recovered into clean tubes, 600 ul of water was added to the polar phase and lyophilized overnight. The apolar phase was dried using a nitrogen downflow for 30 minutes. Both polar and apolar samples were stored dry at -20°C until analysis. Samples were reconstituted either in 200 ul of acetonitrile/water (1:1 v/v) or 100 ul isopropanol/acetonitrile/water (4:3:1, v/v/v) for polar and apolar samples respectively. For aploar samples SPLASH II Lipidomix internal standard was used.

For polar metabolites 1-μl of sample was injected for non-targeted metabolomics analysis by an Ultimate 3000 UHPLC system (Thermo Scientific, Dreieich Germany). Using a binary solvent system (A: 0.1% formic acid in water, B: 0.1% formic acid in acetonitrile) metabolites were separated on a BEH-Amide column (100 mm × 2.1 mm,1.7 μm particle size, Waters, Massachusetts, USA) with a VanGuard 2.1 mm × 5 mm guard column of the same material, by applying a linear gradient from 99% B to 40% B in 6 minutes and then to 4% B in 2 minutes at a flow rate of 0.4 ml/min, returning to initial conditions in 0.1 minute and equilibration for 2 minutes (10 minute total run time). From apolar metabolites 10 μl of sample was injected by an Ultimate 3000 UHPLC system (Thermo Scientific, Dreieich Germany). Using a binary solvent system (C: 0.1% formic acid and 10 mM ammonium formate in 60% acetonitrile 40% water, D: 0.1% formic acid and 10 mM ammonium formate in 90% 2-propanol and 10% acetonitrile) metabolites were separated on a CSH-C18 column (100 mm × 2.1 mm,1.7 μm particle size, Waters, Massachusetts, USA) with a VanGuard 2.1 mm × 5 mm guard column of the same material, by applying a linear gradient from 40% D to 43% D in 3 minutes then to 50% D in 0.1 minutes followed by a change to 54% D in 10 minutes then to 70% D in 0.1 minutes and to 99% D in 5.9 minutes holding at 99% D for 0.5 minutes before returning to initial conditions in 0.1 minutes and equilibration for 1 minute at a flow rate of 0.4 ml/min (20 minute total run time).

Commonly between the two fractions, eluting analytes were electrosprayed in positive and negative mode (two separate injections) by an Apollo II ion funnel ESI source (Bruker Daltonics Inc). Source settings were as follows: capillary voltage 4500 V or -3500 V; end plate offset 500 V; drying temperature 250 °C; desolvation gas (nitrogen) flow 8.0 l min−1; nebulizer gas pressure 3.0 bar. Samples were analyzed by an quadrupole time of flight mass spectrometer (tims TOF Pro, Bruker, Bremen Germany) in tims-off mode, auto MS/MS settings were: switching threshold 350 clear-to-send (cts); cycle time 0.5 s; active exclusion after 3 spectra; release after 0.2 min reconsidering precursors if ratio current/previous intensity > 2. Analytes selected for fragmentation were fragmented by collision with nitrogen gas at a collision energy of 20-30 eV. Precursors and fragments were analyzed by the time-of-flight analyzer using a range of 20-1300 m/z.

The resulting data were analyzed using Metaboscape 5.0 (Bruker Daltonics, Bremen, Germany) to perform data deconvolution, peak-picking, and alignment of m/z features using the T-ReX 3D peak extraction and alignment algorithm. All spectra were recalibrated on an internal lockmass segment (NaFormate clusters) and peaks were extracted with a minimal peak length of 8 spectra (7 for recursive extraction), and an intensity threshold of 1000 cts. The resulting feature tables contain deconvoluted features (adducts combined based on chromatographic correlation and known mass differences). Features were annotated, using SMARTFORMULA (narrow threshold, 1.5 ppm, mSigma:10; wide threshold 5.0 ppm, mSigma:30), to calculate a molecular formula. Spectral libraries including Bruker MetaboBASE 3.0, Bruker HDBM 2.0, MetaboBASE 2.0 in silico, MSDIAL LipidDBs, MoNA VF NPL QTOF, and GNPS, were used for feature annotation (narrow threshold, 1.5 ppm, mSigma 10, msms score 900, wide threshold 5.0 ppm, mSigma:30 msms score 800). Analyte lists of in house measured validated standards were also used to annotate features based on mass, retention time and fragmentation spectrum narrow threshold, 1.5 ppm, RT within 6 seconds, mSigma 10, msms score 900, wide threshold 5.0 ppm, RT within 12 seconds, mSigma:30 msms score 800). An annotated feature was considered of high confidence if more than two green boxes were present in the Annotation Quality column of the program and low confidence if less than two green boxes were present. We also identified metabolites by analysis with SIRIUS (), only retaining structure annotations with a confidence score greater than 0.5.

For further downstream analysis the statistical programming language R (v4.3.3) was used. For both metabolite fractions separately, blank contaminants were removed, when a metabolite was present in the blank samples at 3 times the intensity compared to the analyte samples. Similarly features with high variability in the QC samples were removed if their coefficient of variance was one standard deviation above their overall mean. We further applied a prevalence filter, such that a feature had to be present in at least 25% of the samples. PCA and RDA analysis were performed with packages factoMineR (v2.11) and vegan (v2.6).

All mass spectrometry data are available in the MassIVE repository (https://massive.ucsd.edu/) via the dataset identifier MSV000096030.

### Statistical analysis

Key fermentation parameter data was shown as mean ± standard error of mean (SEM). The significant difference was analyzed by repeated one-way ANOVA using Dunnett post hoc test (GraphPad prism 8). If not indicated differently, signicance levels are set at *P*-value < 0.05.

## Results

### Microbial conversion of MAN to NOR occurs in the colon, not the small intestine

Given the negligible oral bioavailability of MAN^3–5^, we hypothesized that prolonged exposure to the intestinal microbiota, may facilitate its microbial metabolism, potentially influencing microbiome composition and function. To this end, we investigated whether gut bacteria could convert MAN into its aglycone, NOR, using human ileostomy effluent (*n* = 8), a proxy for small intestinal microbiota^19^, and fecal samples from healthy adult volunteers (n = 15). Ileostomy samples were incubated aerobically, while fecal samples were incubated anaerobically, with 500 µM MAN for 48 h. The concentration of MAN was based on a 900 mg oral dose of *M. indica* bark extract (Vimang®)^34–36^, containing 20 wt% MAN^37^, in a colon volume of 819 mL^38^. Samples were analyzed at different time points using High-Performance Liquid Chromatography coupled with UV detection (HPLC-UV).

No MAN conversion was observed in any of the ileostomy samples up to 48 h (**Supplementary Figure 1A**). In contrast, MAN conversion occurred in 6 out of 15 fecal samples after 48 h, but only when enriched beef broth (EBB) without glucose was used as the anaerobic growth medium (**Figure 1B, Supplementary Figure 1B-C)**. The conversion was accompanied by the emergence of a new peak in the HPLC-UV chromatogram, which matched the reference standard of NOR and was confirmed by LC-MS (**Figure 1B**, (**Supplementary Figure 1D**). Among the fecal samples, distinct conversion profiles were observed: 5 samples exhibited complete MAN conversion (High Converters; 84-100% conversion of MAN), 1 sample showed partial conversion (Intermediate Converter; 49% conversion), and 9 samples showed no conversion (Non-Converters) (**Figure 1C**). No additional metabolites were detected, suggesting that the gut microbiota selectively cleaves the C-glycosidic bond of MAN to produce the aglycone NOR. Overall, our findings demonstrate that colonic, but not small intestinal, bacteria mediate the conversion of MAN to NOR. This highlights significant interindividual variation in microbial conversion capacity. However, efforts to culture and isolate the specific bacterial strains responsible for NOR production were unsuccessful.

### Distinct microbiota responses to NOR and MAN supplementation

To investigate the impact of gut bacterial deglycosylation of MAN to NOR on microbiota composition and metabolic activity, we conducted an *ex vivo* fermentation experiment using fecal samples from healthy donors (*n* = 6). Samples were incubated in the SIFR® fermenter platform, consisting of 54 bioreactors, under conditions simulating the colonic environment.^39^ Each sample was treated with MAN, NOR, or no substrate (NSC, control) for 24 hours at 37 °C with an initial pO2 of 15 mmHg (**Figure 2A**). pH, gas production and total cell counts were monitored, and samples were analyzed using shotgun metagenomic analysis, as well as targeted and untargeted metabolomics.

**Figure 2.**
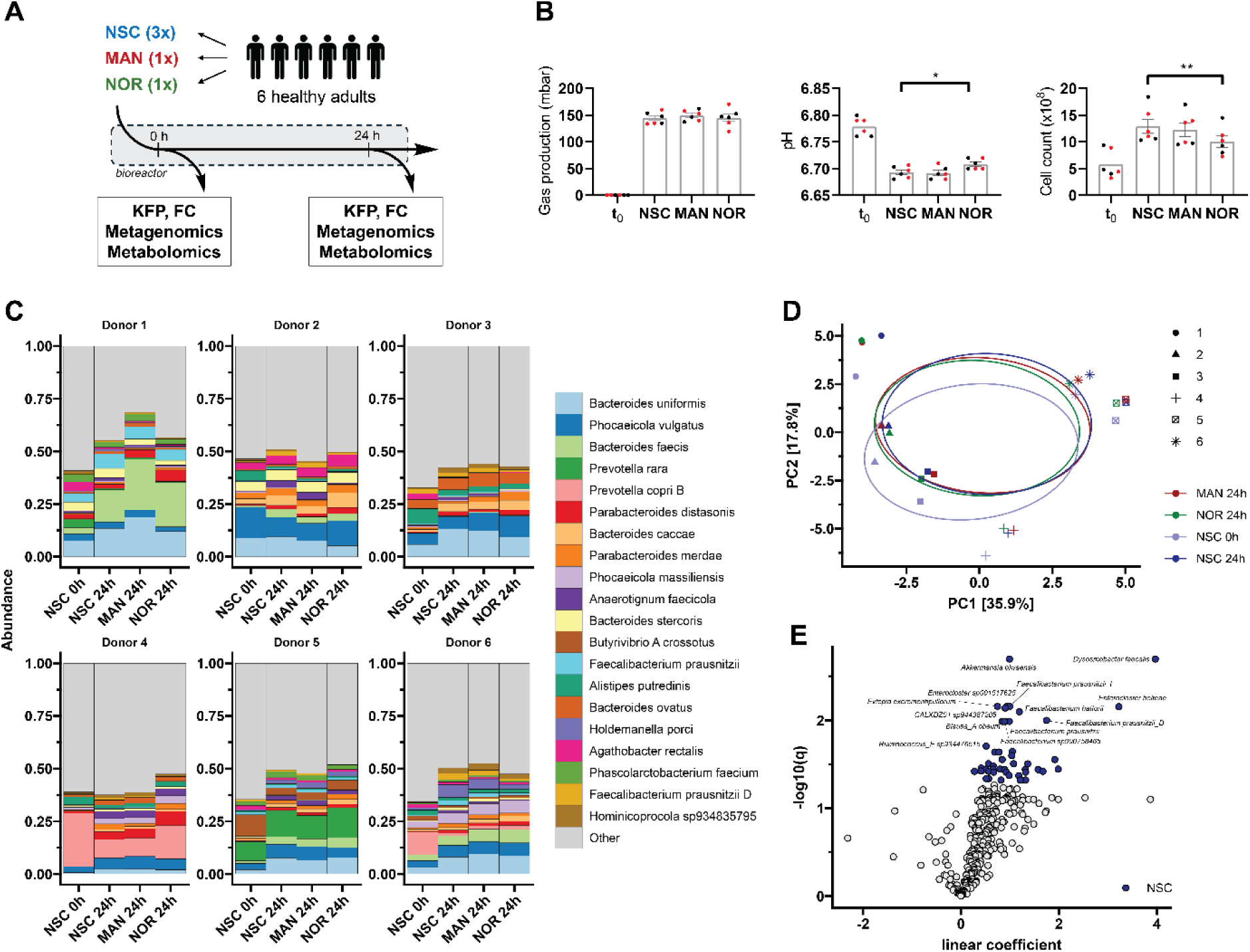
Gut microbiota composition is marginally impacted by MAN or NOR supplementation *in vitro*. (A) Schematic representation of the experimental setup for the fermentation experiment. KFP = key fermentation parameters; FC = flow cytometry; DNA = DNA extraction for metagenomic analysis; HPLC-UV = preparation of samples for targeted metabolomics via HPLC-UV analysis. (B) Bar plots showing key fermentation parameters, such as gas production, pH, and total cell count. Values are expressed as mean ± standard error of the mean (SEM). Statistical significance was assessed using repeated one-way ANOVA with Dunnett’s post hoc test, with significance set at p < 0.05. (C) Microbiota composition showing the top 20 most abundant species per donor and condition. (D) Principal component analysis (PCA) of microbiota composition shows major clustering by donor, with point shapes representing donor ID. Ellipses indicate confidence intervals, and explained variances are shown in brackets next to the axes. (E) Volcano plot of differential abundance analysis using Maaslin2, comparing NOR against NSC at 24 hours. Blue indicates higher abundance in the NSC condition at p < 0.05.

MAN conversion to NOR was confirmed in 3 out of 6 donors, exhibiting moderate conversion rates (13–50%) (**Figure 1**), **Supplementary Figure 2A**). Notably, NOR supplementation significantly altered fermentation parameters, including an increase in pH and a reduction in total cell count (**Figure 2B**), while MAN had no detectable effects on these parameters. Gas production remained unaffected across all treatments. To determine shifts in microbiota composition, we employed deep shotgun metagenomic sequencing. As our aim is to resolve yet uncultured and unknown species which might be able to convert MAN to NOR, we performed *de novo* metagenomic assembly and genome binning with subsequent back-mapping to determine bin abundances. This approach allowed us to resolve up to 355 non-redundant bins per sample after clustering at 95% sequence identity. To overcome statistical challenges in compositional microbiome analysis, relative read counts per bin, expressed through a pseudo TPM (transcripts per million) value, were transformed to absolute abundances using the previously obtained cell counts.

At the community level, samples were mostly composed of *Bacteroides*, *Prevotella*, *Phocaeicola* and *Faecalibacterium* species (**Figure 2C**). The overall composition remained stable between 0h and 24h, but a general increase in Bacteroides species was observed (see **Supplementary Excel Sheet** for differential abundance tests). Microbial richness and diversity did not differ significantly between treatments and control although NOR supplementation showed a tendency to decrease the inverted Simpson index, suggesting a modest reduction in microbial diversity **(Supplementary Figure 2B)**. Principal component analysis (PCA) revealed strong donor-specific clustering, with samples from timepoint 0 h slightly separated along PC2 from those at 24 h **(Figure 2D)**. This pattern highlights consistent abundance shifts across all donors during culturing. However, differential abundance analysis using Maaslin2 revealed marginal effects of MAN supplementation (**Supplementary Figure 2C**), with only *Dysosmobacter faecalis* showing a significant decrease in abundance in MAN treated samples (p < 0.05). In contrast, NOR supplementation led to significant decreases in multiple Clostridia species, including amongst others *Faecalibacterium* sp., *Ruthenibacterium* sp., *Blautia* sp. and *Dorea* sp., which were found to be lower in abundance (**Figure 2E**). Together, these results demonstrate that NOR supplementation, but not MAN, induces distinct alterations in microbiota composition and fermentation parameters. Specifically, NOR reduced the abundance of key Clostridia species and showed a tendency to decrease microbial diversity, emphasizing its selective impact on the colonic microbiota.

### MAN conversion is linked to uncultured species harboring DfgAB genes

Given the broad variability in MAN conversion among the fecal samples and the unsuccessful efforts to culture and isolate the specific bacterial strains responsible for NOR production **(Figure 1)**, we aimed to identify bacterial species potentially responsible for this conversion in our metagenomics data. Differential abundance analysis was performed to correlate conversion percentage with bin abundance estimates. Several species showed significant positive associations with conversion strength (p < 0.05; **Figure 3A**), including *Duodenibacillus* sp900552545, CAKRHR01 sp934339005, and *Phocaeicola plebeius*. While previous studies have identified *Bacteroides* sp. MANG and *Lachnospiraceae* strain CG19-1 as species capable of converting MAN to NOR^10,11^, these were not detected in our samples. To further explore whether the identified species might participate in MAN conversion, we searched for homologs of the DfgAB gene cluster, previously shown to mediate MAN deglycosylation^12,40^, within our non-redundant protein reference catalogue. This analysis revealed the presence of DfgAB genes in four bins corresponding to two species: CAKRHR01 sp934339005 and *Enterenecus* sp900549885.

**Figure 3:**
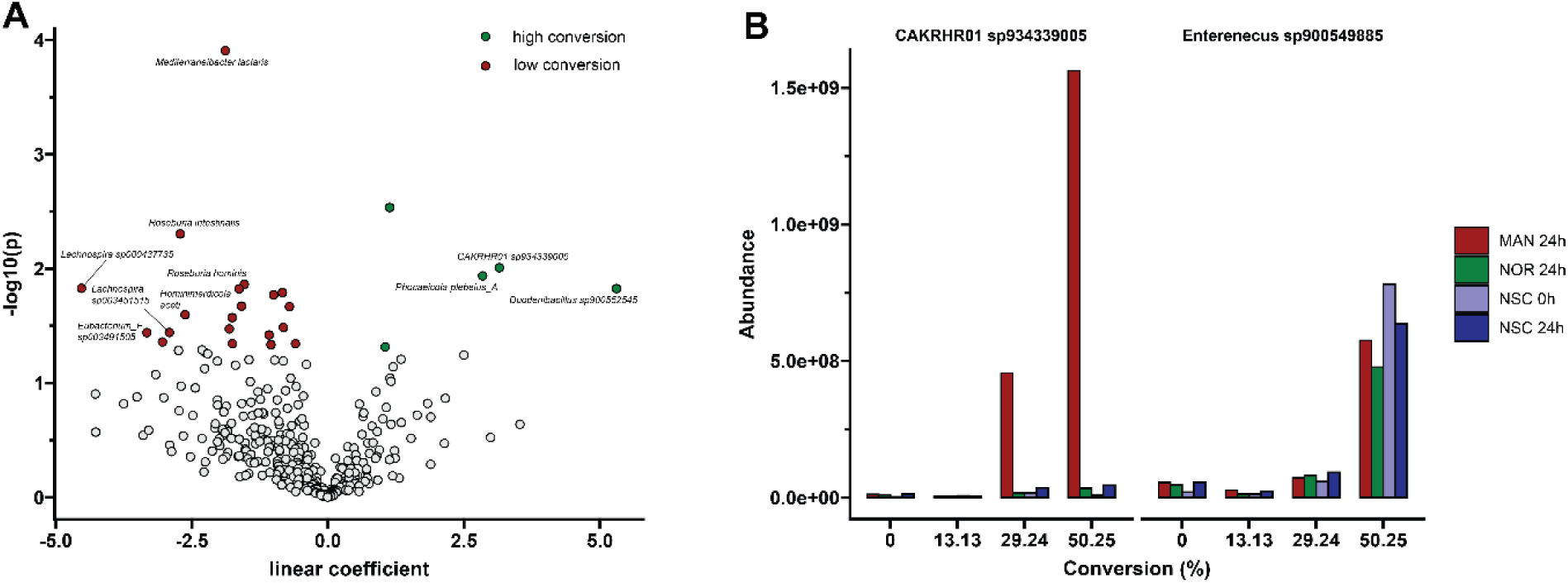
MAN conversion is predicted by microbiota composition. (A) Volcano plot of differential abundance analysis using Maaslin2, associating MAN conversion percentage with bin abundances. Green indicates a positive association with conversion percentage (p < 0.05), while red indicates a negative association. (B) Bar plot showing the abundances of reference bins from species carrying DfgAB homologs across different converter groups.

In *Enterenecus* sp900549885, DfgA and DfgB were located adjacent to one another and surrounded by carbohydrate-processing genes, suggesting their involvement in a broader pathway **(Supplementary Excel Sheet)**. Conversely, in CAKRHR01 sp934339005 the DfgAB genes were dispersed across the genome, with an additional copy of DfgA present. Intriguingly, homologues of DfgCDE were also identified in CAKRHR01 sp934339005, though not organized in a consecutive gene cluster, indicating possible involvement in multiple functions.

Notably, CAKRHR01 sp934339005 displayed a marked increase in abundance in MAN-treated samples that exhibited NOR conversion after 24 h of culturing (**Figure 3B**). In contrast, *Enterenecus* sp900549885 did not respond to the MAN treatment but still showed a positive association with converter status. Together, these findings demonstrate that the presence of DfgAB harboring, yet uncultured, bacterial taxa can predict MAN conversion in an abundance-dependent manner.

### NOR supplementation alters microbial metabolism of SCFA production and urate metabolism

To investigate the metabolic changes associated with the supplementation of MAN or NOR supplementation, short-chain acid (SCFA) levels were measured and samples were subjected to untargeted metabolomic analysis using LC-MS/MS. Except for butyrate, the levels of acetate, propionate, valerate, and bCFAs (sum of isobutyrate, isovalerate, and isocaproate) were significantly reduced by NOR treatment (**Figure 4A**). These changes were also reflected in the total SCFA levels, which was significantly decreased as well in NOR-treated samples. In contrast, MAN supplementation resulted in a significant reduction in overall SCFA levels, particularly propionate and valerate, but with a smaller effect compared to NOR treatment.

**Figure 4:**
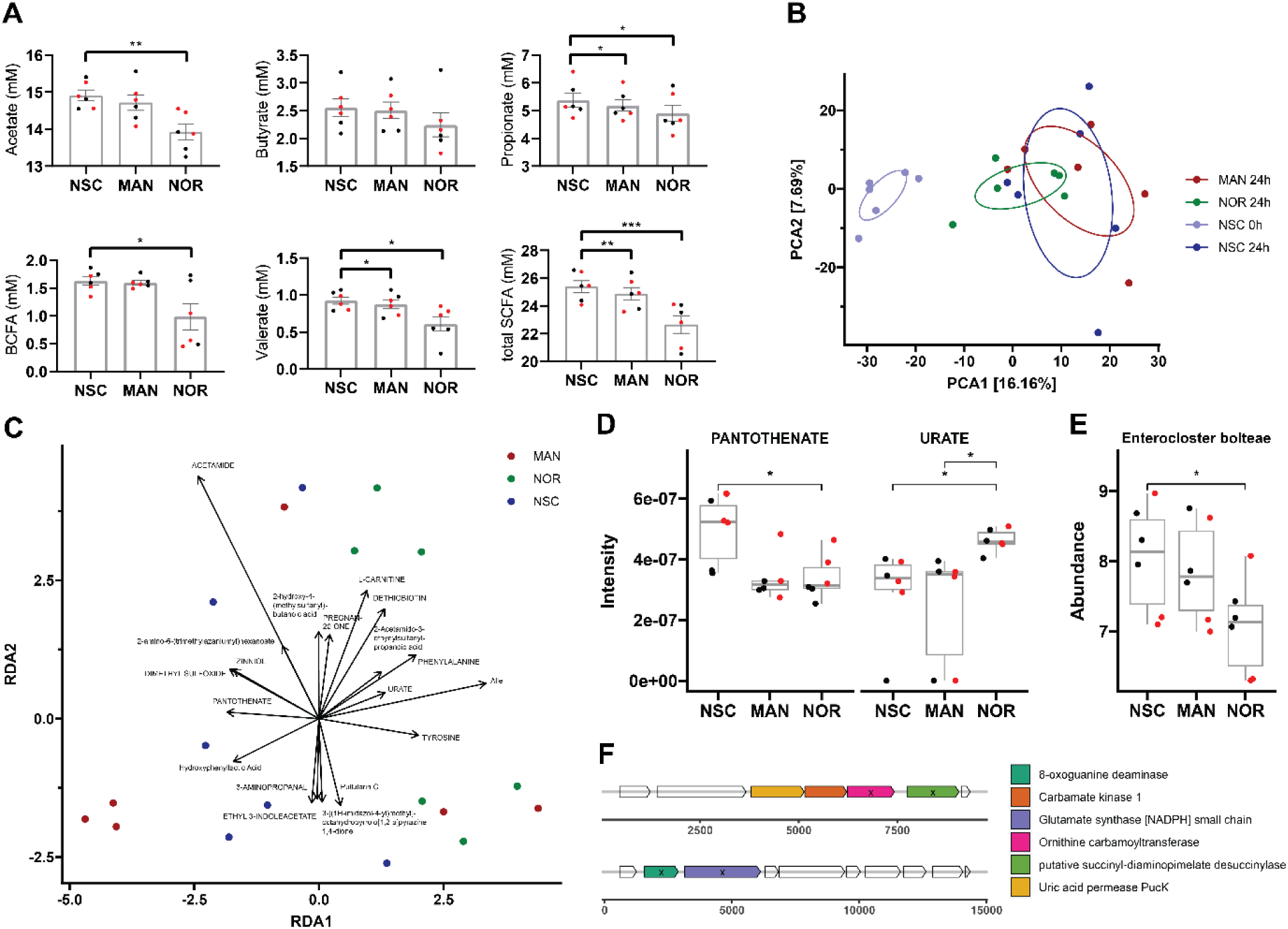
NOR treatment impacts community metabolic activity in vitro. (A) Bar plots showing levels of SCFAs. Data points in red represent samples from donors with MAN conversion activity. Values are expressed as mean ± standard error of the mean (SEM). Statistical significance was determined using repeated one-way ANOVA with Dunnett’s post hoc test (p < 0.05). (B) Principal component analysis (PCA) showing major clustering of inoculum and NOR-treated samples. Confidence ellipses are shown, with explained variances indicated in brackets next to the axes. (C) Redundancy analysis (RDA) reveals grouping of NOR-treated samples. Arrows represent the top 20 metabolites with the strongest associations to each group, with arrow length proportional to the strength of the association. Metabolites in capital letters are identified by spectral matching, while others are identified by SIRIUS. (D) Mass spectrometry intensities of polar metabolites identified from the RDA, showing significant differences in NOR-treated samples (t-test, p < 0.05). (E) Absolute abundance of *Enterocloster bolteae* shows a significant reduction in NOR-treated samples (t-test, p < 0.05). Data points in red represent samples from donors with MAN conversion activity. (F) Genetic arrangement of genes involved in urate metabolism, identified by homology (denoted with ‘x’).

To gain deeper insight into the broader metabolic changes, we performed untargeted metabolomics from both polar and apolar metabolite fractions, separately. Metabolites were filtered for blank contamination and QC variability and identified by spectral matching. PCA analysis revealed distinct clustering of treatments for both fractions, especially the polar one (**Figure 4B; Supplementary Figure 3A**). Interestingly, while inter-donor variation in microbial composition was prominent (**Figure 2D**), the metabolomics data showed significantly less variation between donors. Samples from the initial inoculum (timepoint 0h) were significantly different (PERMANOVA, p<0.002) from samples after fermentation. While no significant differences in clusters were observed after 24 hours of incubation, NOR samples exhibited lower dispersion (average distance to centroid: 21.91 vs. 23.01 [MAN], 24.21 [NSC]) and formed slightly distinct clusters along PCA1 compared to MAN and NSC. This trend was further supported by RDA analysis, constrained to treatment and adjusted for residual donor variation (**Figure 4C**).

Notably, two MAN-treated samples from donors exhibiting converter activity grouped with NOR-treated samples, indicating similar metabolic shifts due to the release of NOR via enzymatic conversion (**Supplementary Figure 3B**). We find that NOR treatment was associated with a higher abundance of amino acids such as phenylalanine or tyrosine as well as urate, L-carnitine and dethiobiotin (**Figure 4C**). Upon closer investigation, pantothenate was significantly reduced in MAN and NOR-treated samples, while urate was significantly increased in NOR-treated samples compared to both MAN and NSC (**Figure 4D**). Other selected metabolites did not show any prominent changes (**Supplementary Figure 3C**).

Previous studies have shown that urate metabolism is prevalent amongst gut bacteria and is governed by a specific gene cluster consisting of up to 7 genes.^41^ To investigate whether our observations coincide with abundance changes in species involved in urate metabolism, we performed homology search against our metagenomic protein catalogue and identified 8 species with 4 to 5 urate cluster genes present, many of which are known to participate in urate breakdown **(Supplementary Excel Sheet)**. Among these, only *Enterocloster bolteae* showed a significant reduction in abundance upon NOR treatment (**Figure 4E**; **Figure 2E; Supplementary Figure 3D**). To confirm the possibility of urate breakdown in this species, we analyzed the genomic context of the four genes. Due to the fragmented nature of its assembly, the urate gene cluster was split across two contigs, however, the genes were in close proximity and an additional uric acid permease was found upstream (**Figure 4F**). Overall, these results indicate that NOR treatment impacts the metabolic activity of the community, leading to reduced SCFA production and alterations in urate metabolism, which are linked to a decrease in the absolute abundance of urate metabolizing species.

## Discussion

In this study, we have demonstrated a moderate prevalence of gut bacteria capable of converting MAN into its aglycone NOR **(Figure 1)**. Our metagenomic analysis identified two yet uncultured strains, *Enterenecus sp900549885* and CAKRHR01 *sp934339005*, which carry homologs of the DfgAB gene cluster implicated in MAN metabolism. Notably, the abundance of CAKRHR01 *sp934339005* was positively correlated with MAN conversion **(Figure 3)**. Finally, NOR treatment led to reduced SCFA production and alterations in urate metabolism, reflecting a shift in metabolic activity linked to a decrease in the abundance of urate-metabolizing species **(Figure 4)**.

Notably, fecal conversion of MAN to NOR was only observed in the absence of carbohydrates in the growth medium, which is consistent with previous reports indicating that C-glycoside conversion is stimulated in carbohydrate-free conditions.^42,43^ These findings suggest that it may be necessary for certain bacterial genes to be activated in low-nutrient conditions before MAN metabolism can occur.

The remarkable cleavage of the C-glycosidic bond in MAN has been previously observed, leading to the successful isolation of two strains from human fecal samples, that were responsible for production of NOR: *Bacteroides sp. MANG* and *Lachnospiraceae strain CG19-1*.^10,11^ In the study describing *Lachnospiraceae strain CG19-1*, the strain originated from 1 out of 19 tested fecal samples which was found to be capable of converting MAN into NOR.^11^ Although the total number of tested samples was not reported for *Bacteroides sp. MANG*, the strain was isolated from a single fecal sample.^10^ These findings suggest that these strains are present in very low abundance among the general human population, which could explain why they were not detected in our study, further emphasizing the interindividual variation in microbial conversion capabilities. Instead, when we correlated our metagenomic data with conversion percentages from targeted metabolomics, we identified, for the first time to our knowledge, two uncultured strains (*CAKRHR01 sp934339005* and *Enterococcus sp900549885*) that are likely involved in MAN deglycosylation. Notably, CAKRHR01 was recently found to share remote sequence similarity (∼60% amino acid identity) with species from the *Catenibacillus* genus which are known to exhibit prominent enzymatic activity against C-glycosides.^44^

Remarkably, NOR supplementation significantly reduced both SCFA and bSCFA production compared to the control, with MAN exerting a similar but less pronounced effect. Although polyphenols are commonly associated with increased SCFA production and enhanced microbial diversity^45,46^, several studies have reported inhibitory effects of isolated polyphenols or polyphenol-rich extracts on specific gut bacterial strains, occasionally accompanied by reduced SCFA levels.^47–51^ Additionally, xanthones have been implicated in promoting microbial imbalance.^52,53^ These findings collectively suggest that certain polyphenols, including xanthones, may selectively inhibit the growth of specific gut microbial members through mechanisms yet to be elucidated, resulting in reduced SCFA production. Consistent with this, our metagenomic analysis revealed a notable reduction in SCFA producing bacteria, including *Faecalibacterium* species, particularly *F. prausnitzii*, underscoring the selective impact of NOR on beneficial microbes.

Furthermore, NOR treatment resulted in a reduction of the beneficial urate-metabolizing strain *Enterocloster bolteae*, which was correlated with a notable increase in urate concentration. Urate-metabolizing commensal strains are vital for modulating systemic uric acid levels, compensating for the absence of uricase in humans by converting urate into acetate and other SCFAs.^41,54^ This aligns with our findings, where increased urate levels were accompanied by a significant reduction in acetate production. These results suggest that NOR may disrupt the balance of urate metabolism, potentially inhibiting SCFA production through the downregulation of urate-consuming bacteria such as *Enterocloster bolteae*. While NOR has been shown in animal models to reduce systemic urate levels by inhibiting its production from xanthine and promoting renal excretion^55,56^, our findings reveal a potentially counterproductive effect on urate metabolism within the gut. This underscores the need for a deeper understanding of how NOR impacts both systemic and gut-specific urate pathways.

Together, these findings indicate that the observed metabolic shifts result from a possible inhibitory effect of NOR on specific Clostridia strains, leading to a reorientation of metabolic activity, as evidenced by the significant changes in urate and SCFA metabolism in NOR-treated samples.

Overall, our study provides new insights into the microbial metabolism of MAN, a compound with a longstanding history in traditional medicine **(Figure 5)**.^1,2^ While MAN itself exhibited limited metabolic effects, its microbial conversion to the aglycone NOR, potentially mediated by specific gut bacteria such as CAKRHR01 *sp934339005* and *Enterococcus sp900549885*, may explain some of the bioactive properties attributed to MAN. The observed metabolic shifts: reduced SCFA production, altered urate metabolism, and selective impacts on key bacterial species, suggest that NOR supplementation could influence gut microbial activity in ways that extend beyond the effects of MAN alone. Additionally, these results urge the need for caution when interpreting the health implications of phytochemicals, particularly for individuals with dysbiosis or those prone to metabolic or gastrointestinal disorders, as well as hyperuricemia. The findings advocate for a personalized approach to dietary and therapeutic recommendations and underscores the importance of understanding the role of the gut microbiome in shaping the pharmacodynamics of such compounds.

**Figure 5:**
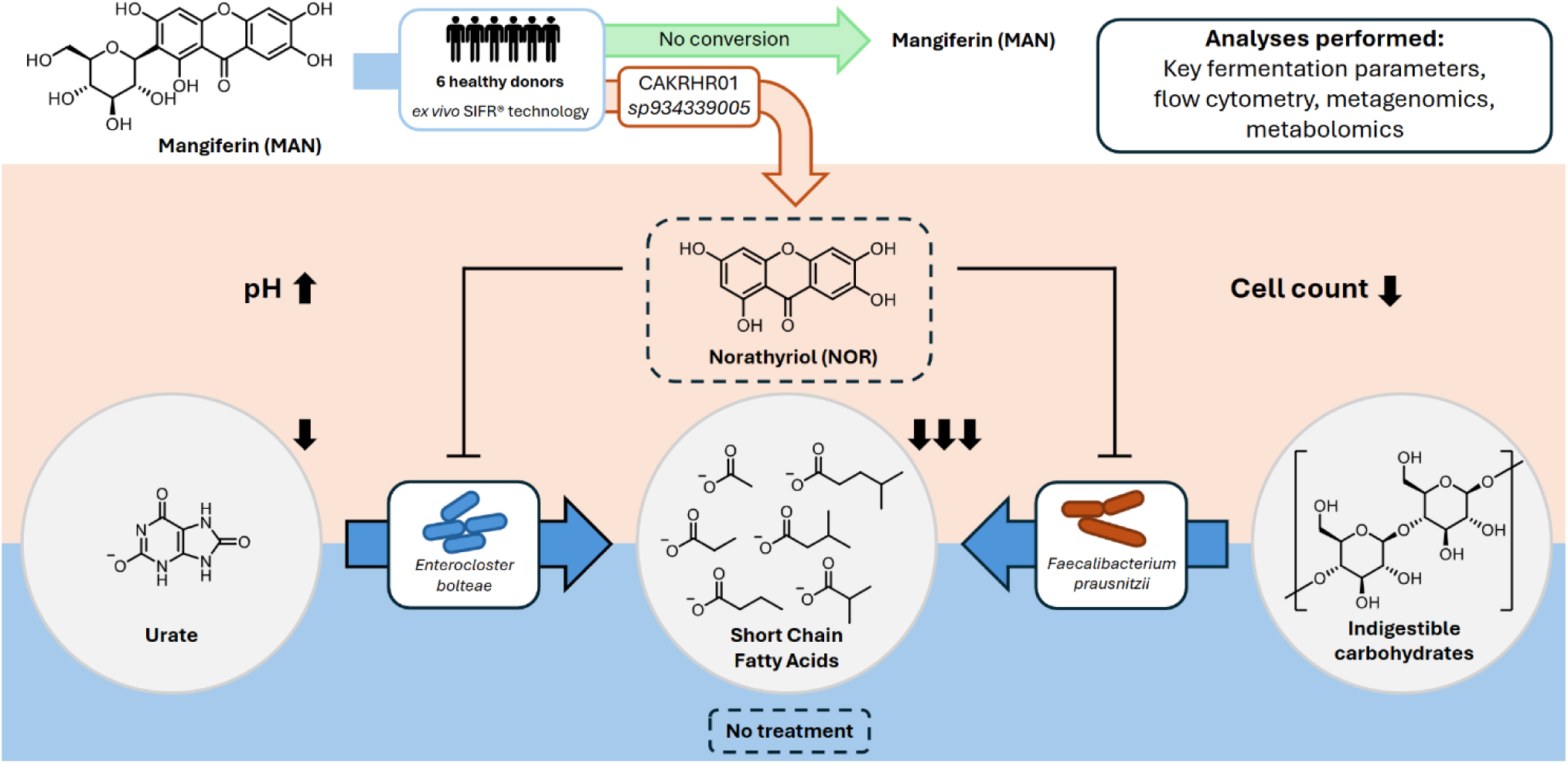
**Overview of the results presented in this study.**

## Acknowledgment

The authors gratefully acknowledge S.P. Kok for establishing a reproducible MAN conversion assay and conducting HPLC-UV analysis.

## Author contribution

DVB, MCS, SEA: Conceptualization, writing-original draft, and editing; DVB, MCS, FH: conducting research; MCS, DVB: data analysis, methodology, preparing figures.

